# On the molecular filters in cochlea transduction

**DOI:** 10.1101/374363

**Authors:** Valeri Goussev

## Abstract

The article is devoted to the specific consideration of the cochlear transduction for the low level sound intensities, which correspond to the regions near the perception threshold. The basic cochlea mechanics is extended by the new concept of the molecular filters, which allows us to discuss the transduction mechanism on the molecular level in the space-time domain. The molecular filters are supposed to be built on the set of the stereocilia of every inner hair cell. It is hypothesized that the molecular filters are the sensors in the feedback loop, which includes also outer hair cells along with the tectorial membrane and uses the zero compensation method to evaluate the traveling wave shape on the basilar membrane. Besides the compensation, the feedback loop, being spatially distributed along the cochlea, takes control over the tectorial membrane strain field generated by the outer hair cells, and implements it as the mechanism for the automatic gain control in the sound transduction.

## Introduction

The cochlear transduction has a long history of research, beginning from the pioneer works of Bekesy [1] and many others contemporary researchers [2-5]. The widely accepted point of view in question is that the mechanical sound vibrations are transformed by the cochlea to the traveling waves of the basilar membrane (BM), which in turn transfers the waves to the outer and inner hair cells. The inner hair cells (IHC), being mechanically bent, open their ion channels, what results in the cell depolarization and subsequent afferents excitation [6]. To be able to explain the experimentally observed high sensitivity of the cochlear transduction for the low level sound intensities an additional feed-forward mechanism for the outer hair cells (OHC) was proposed, which actively amplifies the basilar membrane vibrations. The whole set of proposed mechanisms was satisfactory in explaining the experimental psychoacoustics data. However the small doubts still exist when remembering the Bekesy’s comment [7] that at 3 kHz “we can hear a vibration of the air particles that is 100 times smaller than the diameter of the orbit travelled by an electron around the nucleus of a hydrogen molecule”. The numerical simulations of the traveling waves for the real sound intensities also confirm the low values of the basilar membrane deformations [8]. It seems pretty unrealistic to assume that the BM displacements of order 0.1-10 nm could cause bending of stereocilia in OHC and IHC strong enough to mechanically open their ion channels. Instead considering the mechanical strips – tip links of the size of a hydrogen atom, which pull mechanically the small ion doors while bending stereocilia, we will try to consider the molecular mechanisms based on the endolymph molecules. They don’t presume any stereocilia bending and tip links and even additional mechanical amplifiers of the basilar membrane traveling waves. The direct participation of the endolymph molecules in the transduction process was discussed a long time ago [9] and then later [10]. We are building upon this idea, providing it with a number of the new views, which being combined together, lead us to the new concept of the space-time molecular filters in the cochlear transduction.

## Model

### Basic mechanics

We consider the basilar membrane deformations based on solution of the conventional set of the hydrodynamics equations describing these deformations as the response to the input sound pressure in 2D uncoiled model of cochlea [2, 11]. The important point for our consideration is to assume that for low level sound intensities the basilar membrane has negligible deformations in the region of the IHC [11]. The OHC, being always in the contact with the tectorial membrane (TM), transfer vibrations of BM to the surface of TM. These vibrations propagate equally in longitudinal and the radial directions and induce the traveling waves in TM. Another assumption is that the vibrations are assumed to be a sum of two independent sources. One of them is passive, evoked by BM displacements. The second one is active and is evoked by OHC via efferent fibers of the cochlear nerve. In contrast with the BM traveling waves, observed basically in the region of OHC rows [11], those of TM are present at all points of its body, except the line of its attachment at the spiral limbus. Hence, the consequence of our assumptions results in the conclusion that IHC are almost immobile and the TM surface in the close vicinity of IHC is submitted to vibrations, which change the distance between TM itself and the top of the stereocilia of every IHC.

The gap between the top of the stereocilia and the TM surface is small and can change essentially during transduction. As estimated from micrograph sources [12] it can be about a size of diameter of the stereocilia, which is about 0.25 mkm [13], i.e. less than the mean height of IHC stereocilia (2-8 mkm [13]). The gap looks like a cave or tunnel of the TM length and is restricted from one side by the Hensen’s stripe.

### The endolymph flow

The movable front of the TM traveling wave initiates locally the endolymph flow in the narrow space between the row of the stereocilia tops and the TM surface. The flow has something in common with the shallow waves [14]. We consider the simplified model of the flow, replacing the tunnel by the 2D space (*x*, *y*), where x coordinate represents the longitudinal direction along IHC row, and *y* coordinate is directed outside from the tops of IHCs in the transverse direction. We limit the further consideration to the TM wave shape *η*(*x,t*), which is locally represented by the pure cosine wave:

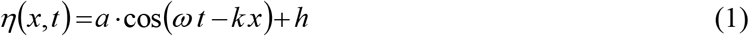

where *a* - the wave amplitude, *ω* - the wave angular frequency, *t* - time, *k* - the wave number, *h* - the tunnel height (*a* < *h*). The amplitude *a* is essentially less than the wave length *λ*, so the wave looks like a flat wave and its space propagation is defined by the parameter *k*. For the travelling wave of 1 kHz frequency and the travel speed about 2 m/s we have *λ* = 2000*mkm*.

The velocities of the flow *u*(*x, y, t*) and *v*(*x, y, t*) can be described by the solutions of the Navier-Stokes (NS) equations separately for the longitudinal *x* and transverse *y* directions. We make some assumptions to simplify the solution of NS equations. As we limit our consideration to the weak sound signals near the perception threshold level, we consider the linearized NS equations. Moreover, considering the time constant 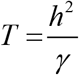 of the transient response of the NS equation solution (*γ* is the kinematic viscosity), we can estimate it for the typical parameters of the flow in the tunnel space *h* = 3 *mkm*, *γ*=1.004· 10^6^*mkm*^2^/*s* [15] as *T* =10 ^5^*s*. Hence, for the real sound signals frequencies up to 20 KHz the transient responses are almost non inertial and all time-dependent events in the tunnel space are simply the transitions between the steady state solutions of the NS equations. Taking into account the made simplifications we obtain the full set of NS equations for our problem as:

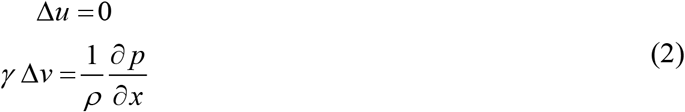

where Δ - Laplace operator, *γ* - the kinematic viscosity, *ρ* - the endolymph density, *p* -the transverse pressure evoked by the TM wave.

The continuity equation

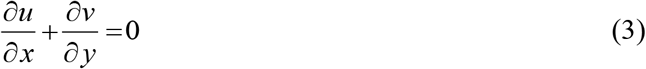

The boundary conditions

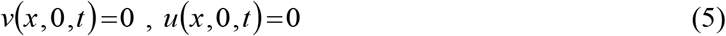

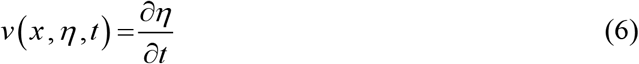

The solutions for velocities *u, v* are given by expressions:

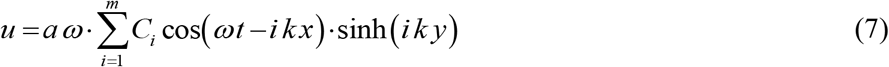

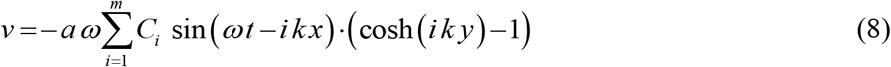

where *C_i_* are arbitrary constants, *m* is the number of the eigenfunctions used for the approximation of *u*(*x, y, t*) and *v*(*x, y, t*). The constants *C_i_* can be evaluated from the boundary condition (6), which looks like the approximation of 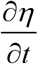 by the series of the eigenfunctions 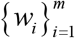:

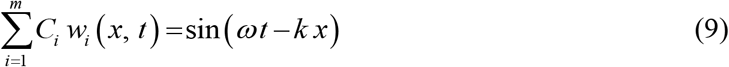

where *w_i_*(*x,t*) = sin(*ωt-ikx*)·((cosh(*ik*·(*a*ηcos(*ωt*–*kx*)+*h*))-1)

In spite that the regular traveling waves on BM have amplitudes in the range 10-50 nm [4], we consider an illustrating example with the amplitude 250 nm, which help us to demonstrate in the more clear way the properties of the endolymph flow inside the IHC tunnel. The obtained results will be valid also for small amplitude ranges and will be used later when describing the molecular movement near the IHC tops.

The velocities *u, v* in *x, y* space build the flow vector field, which is represented for *a* = 0.25*mkm*, *h* = 0.75*mkm*, and m=5 on the Fig.2:

**Figure.1.**
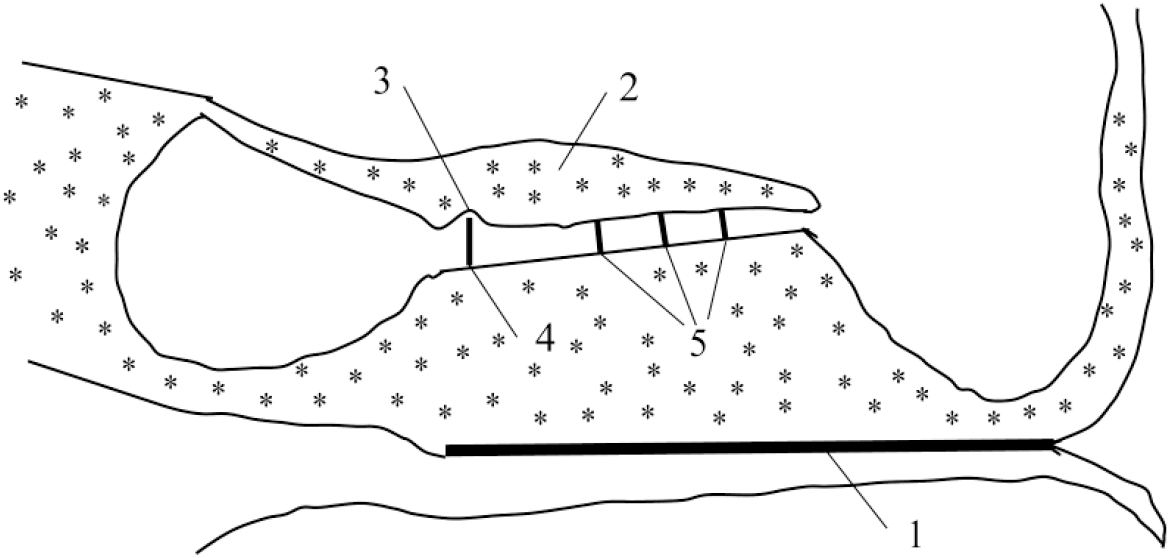
Schematic representation of the tectorial subspace near IHC region, 1 -the basilar membrane (BM), 2-the tectorial membrane (TM), 3 – the IHC tunnel, 4 – the inner hair cells (IHC), 5 - the outer hair cells (OHC).

**Figure.2.**
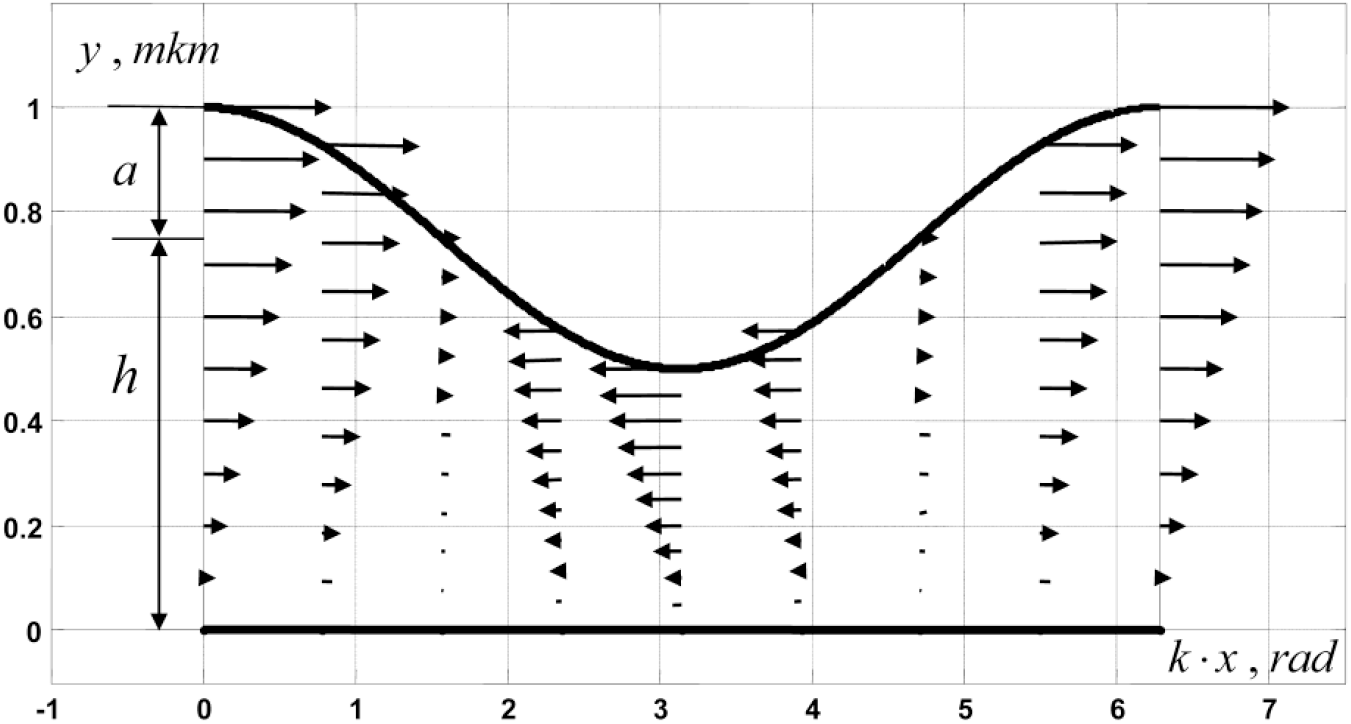
The vector field of the flow in the IHC tunnel, (the velocity module is down scaled by the factor 5.15 ·10^-7^)

We can see that the values of the transverse velocity *v* are almost negligible as compared with the longitudinal velocity, what is typical for all shallow waves [14]. We define the instantaneous angular velocity Ω(*x,t*) of the flow in every point of the wave *η*(*x,t*) relative to the IHC tops:

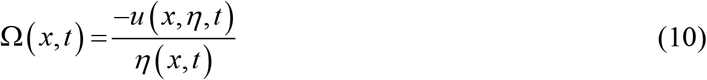

The most essential property of the tunnel longitudinal flow is the dependence of its angular velocity Ω(*x,t*) on the tunnel height *h* (Fig.3). Decreasing the height may serve amplifying the angular flow velocities for weak sound signals. The maximal module of the angular velocity is associated with the backward propagation of the traveling wave, as it is closer in this region to the IHC tops.

**Figure.3.**
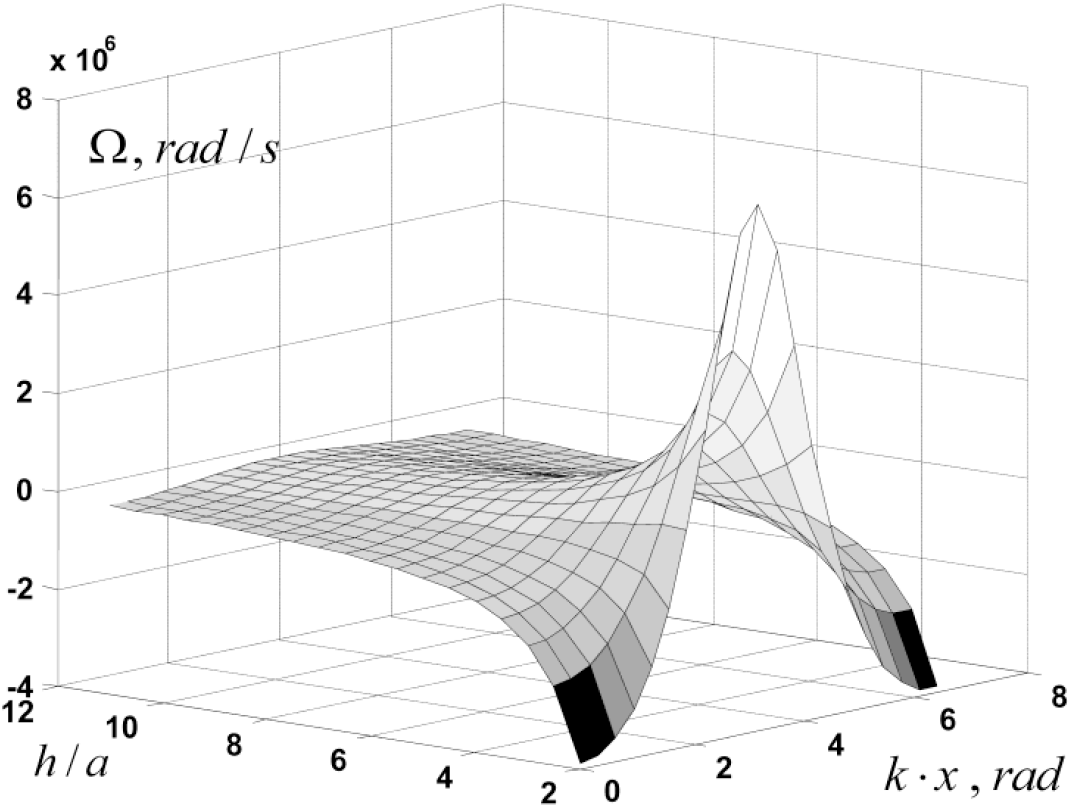
The gain of the tunnel flow angular velocity as a function of the tunnel height

### Molecular sensors

According to the early published works [10, 16-19] the endolymph consists of the acid mucopolysaccharide molecules, which are considered as the long threads of the length about 0.5 mkm and the width about 4 nm. One of them most imported features is that they are getting polarized, being bent by the hits of surrounding water molecules submitted to the Brownian motion. The main assumption of our approach is that the connection of any two stereocilia tops by the polarized endolymph molecule is able to open the ion channels in both stereocilia. The length of the endolymph molecule surprisingly fits the doubled mean stereocilia diameter. The cumulative effect of such connections along the whole set of stereocilia in every IHC can lead to the cell depolarization and the afferent neuron activity. The same approach was early proposed for the explanation of the vestibular transduction [20]. The endolymph molecules are moving in the complex environment, being submitted to Brownian movement and also the tunnel flow. That’s why the connection of the molecule with the stereocilia tops is the stochastic event and needs to be considered in more detailed way. Let us consider the endolymph molecule as a thin rigid beam (Fig.4).

**Figure.4.**
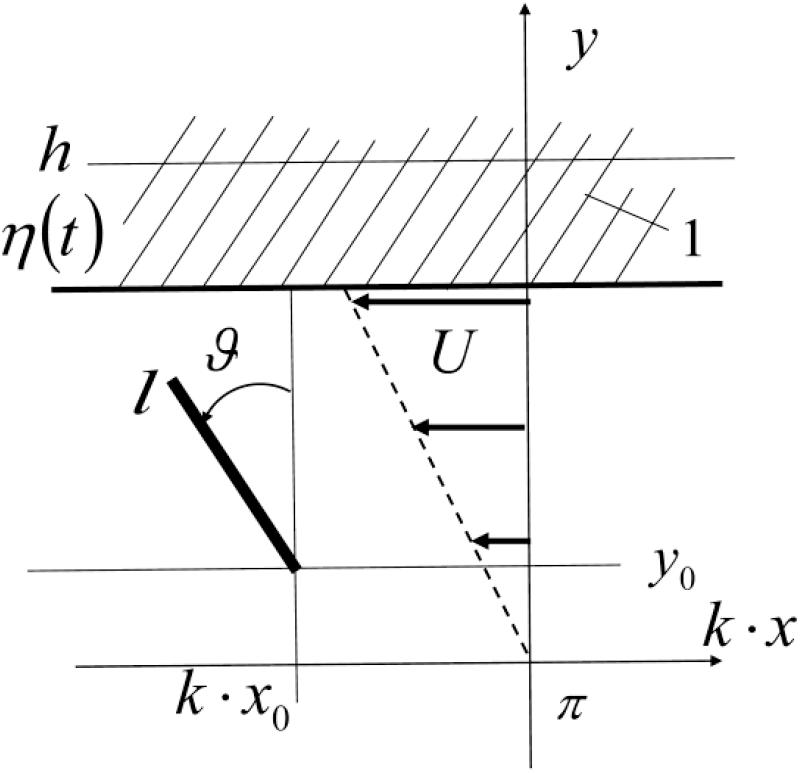
Movement of the endolymph molecule in the IHC tunnel evoked by the TM traveling wave, 1-the fragment of the tectorial membrane, U – boundary velocity, *η*(*t*) - deflection of the tectorial membrane surface from the IHC top position, l – length of the endolymph molecule, *ϑ* - its angular position.

The line *y* = 0 will be associated with the stereocilia tops. The line *y* = *η* represents the local tunnel height at *k* · *x* = *π*, where the maximal longitudinal flow velocity *U* is observed. Considering the small deflections from the IHC tops, we linearize the longitudinal velocity *u* in the variable *y*, so it will be similar to the Couette flow. After the simplification the velocity *U* reads: 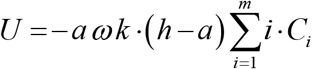. Every endolymph molecule in space *x, y* is described by its position point *x*_0_, *y*_0_ and the orientation angle *ϑ* counting from *y* axis. All points *x*_0_, *y*_0_ and angle *ϑ* are randomly distributed inside the space. The molecule is supposed to be able to make evolution with the angle *ϑ* greater then 2*π*, *ϑ* ∈ [−∞, ∞]. All endolymph molecules independent of their position are affected by the same angular velocity 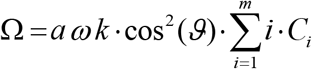 of the drift flow. The maximal value Ω_*m*_ of Ω is reached at *ϑ*=0 :

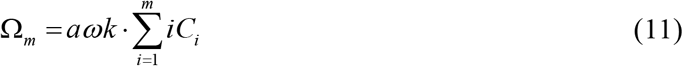

The endolymph molecules are submitted to the Brownian motion and the drift and their stochastic behavior can be described by the solution of the Fokker-Plank equation for the probability density function for the molecule orientation [21] as:

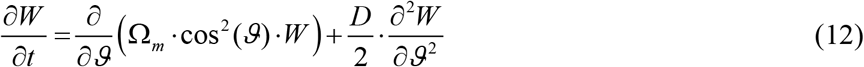

where *W* = *W*(*ϑ, t*) is the probability density function, *D* is the molecule diffusion coefficient in rotation. The solution of Eq (12) has two components: one is defined by the rotational diffusion, and another related to the drift by the Couette flow velocity. As the diffusion is related only to the angle distribution about the mean value, we are more interesting in the mean angle changes evoked by the Couette drift velocity. To get them we multiply Eq (12) by *ϑ* and integrate it in the angle space [−∞, ∞]. Integrating by parts and removing the null components, we finally obtain:

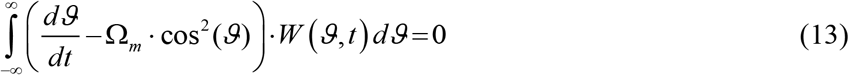

As the distribution function *W*(*ϑ, t*) independently changes in time, the only way to get Eq(13) valid is to assume that:

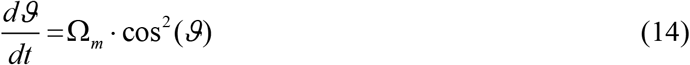

Equation (14) should be valid for all values *ϑ* in the distribution *W*(*ϑ, t*) and describes the drift trajectory for the particular value 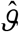 taken from *W*(*ϑ, t*). Integrating equation (14) in time yields:

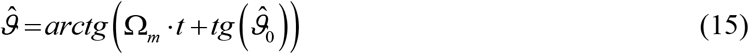

where *t* is the observation time, 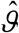 is the drift evoked component of *ϑ* at the time *t*, 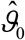 is the initial drift angle. The initial angle 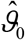 is an arbitrary angle taken from *W*(ϑ,0) distribution. We assume that in the absence of any drift flow the initial orientation angles of the endolymph molecules are distributed uniformly in the range 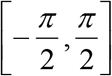 with the probability density function 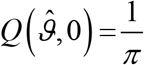. From equation (15) we can see that the drift changes the initial distribution of 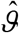 in time. The changed distribution 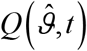 reads:

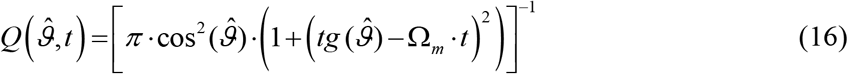

We can see that 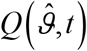 changes gradually from the uniform distribution at Ω_*m*_ = 0 to the δ-function distribution at Ω_*m*_ =œ. As it is reported [22] approximately constant traveling wave velocity at the definite point of BM for wide range of the sound intensities, we can conclude that only the local tunnel height *η* is able to change Ω_*m*_ and the shape of the density function. Hence, the amplitude of the TM traveling wave, which changes the tunnel height locally, changes in the same time the shape of the probability density function 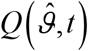. We can also say that all information about the incoming sound amplitude and phase, which is available in the TM traveling wave, is reflected also on the density function 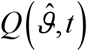, i.e. by the endolymph molecules orientation in every point of the tunnel longitudinal axis.

The set of particular density functions 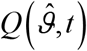 is represented on Fig.5 for 7 height values, the traveling wave amplitude *a* =10*nm*, the wave frequency 1 kHz, and the observation time *t* = 50*mks*.

**Figure.5.**
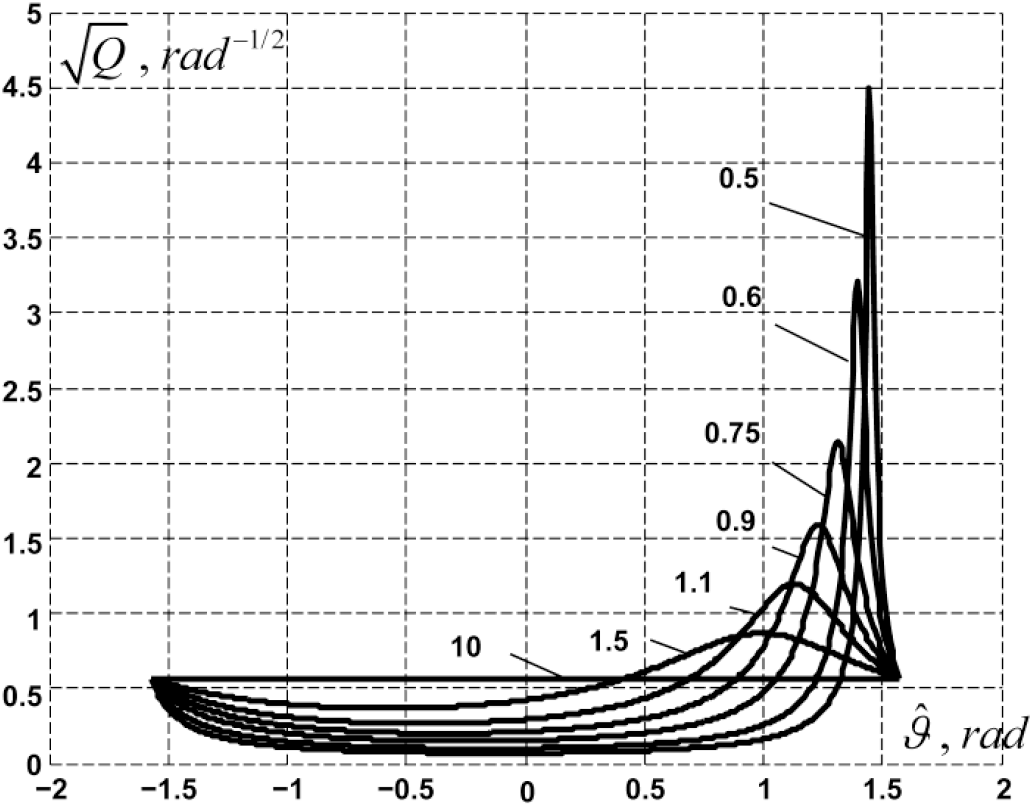
The probability density function 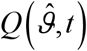 for various heights *h*(*mkm*) and *a* =10*nm*.

For better visibility all density functions were transformed using the square root function. The Fig.5 is an illustration of the basic idea that to get IHC depolarized the endolymph molecule orientation should be close enough to the line of the stereocilia tops. Let us define the angle *β* ∈[0, *π*] counting from the line of stereocilia tops 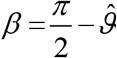. From the density function 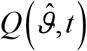 we can evaluate the probability of the molecule orientation to be in the close vicinity to the tops line :

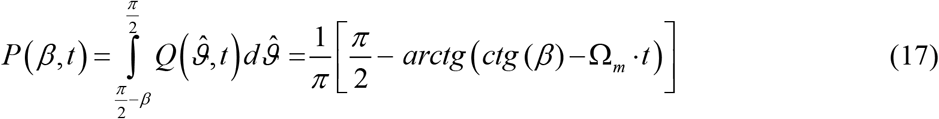

The set of the probability curves, corresponding to the particular density functions Fig.5 is represented on Fig.6.

**Figure.6.**
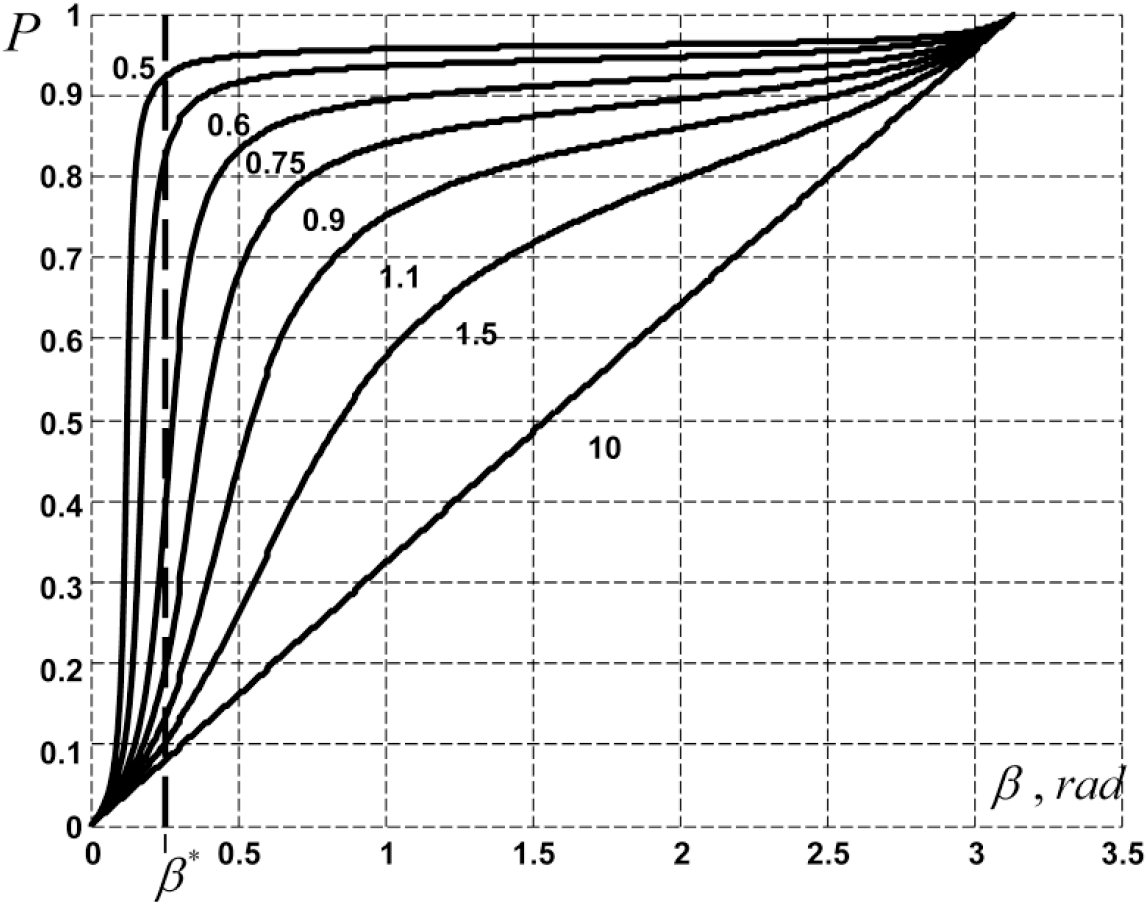
The probability of IHC excitation for different tunnel heights *h*(*mkm*) and *a* =10*nm*

Let us define the threshold *β*\* of the molecule orientation and the depolarization region *β* ≤ *β*\*, inside which the IHC can be depolarized. Then the corresponding probability *p*\* = *P*(*β*\*, *t*) will give us the probability of depolarization at the boundary of the depolarization region. It is then obvious that the high level of the IHC probability depolarization corresponds to the high level of the spike activity in the afferent neurons. All probability curves presented on the Fig.6 are related to the separate transduction event occurring in vicinity of one pair of IHC stereocilia. In reality, every IHC has about 5-10 pairs of stereocilia, separated by the endolymph molecule length. Assuming that the particular probabilities of every pair of stereocilia are independent, the total probability excitation for one IHC *P_ex_* will be the product of the particular probabilities *P_i_* (*β*\*, *t*) of every pair of stereocilia.

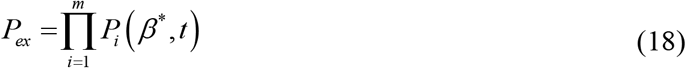

where *m* is the number of the stereocilia pairs in IHC.

Defining the gain *δ* of the IHC in detection of the weak sound signals as the quotient of two probabilities of excitation *P*_1_ to *P*_2_ for the tunnel heights *h*_1_ and *h*_2_ as:

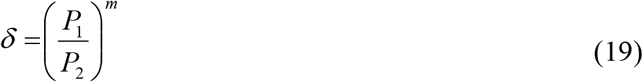

we evaluate from Fig.6 for *h*_1_ = 0.5*mkm*, *h*_2_ = 10*mkm*, and *m* = 5 the gain as *δ* ≈ 5.9 ·10^4^.

The probability of the IHC depolarization and the activity of the afferent neurons are depending on the height of the gap between the stereocilia tops and the TM, and on the traveling wave amplitude. The gain *δ*, which defines the sensitivity of the IHC molecular filter, can be changed by the auditory cortex using the low frequency depolarization potentials in OHC signal innervations, which will make changes in the static length of OHC.

### Space-time filtering

The specific properties of the endolymph flow in the tunnel and those of the molecular filters are supposed to be integrated in the general feedback control loop, which consists of all IHC, OHC sets, BM,TM, and the auditory cortex. The control loop has distributed parameters at least along the longitudinal axis and time depended properties. Its main task can be the best estimation of the BM vibrations in presence of noise and disturbing harmonic signals, so it can be considered as the space-time filter. Let us consider the general filter structure (Fig.7), which is intended to reproduce the input signal *s*(*x, t*) (the TM vibrations evoked by the BM and distributed along the longitudinal axis *x* of the uncoiled cochlea).

**Figure.7.**
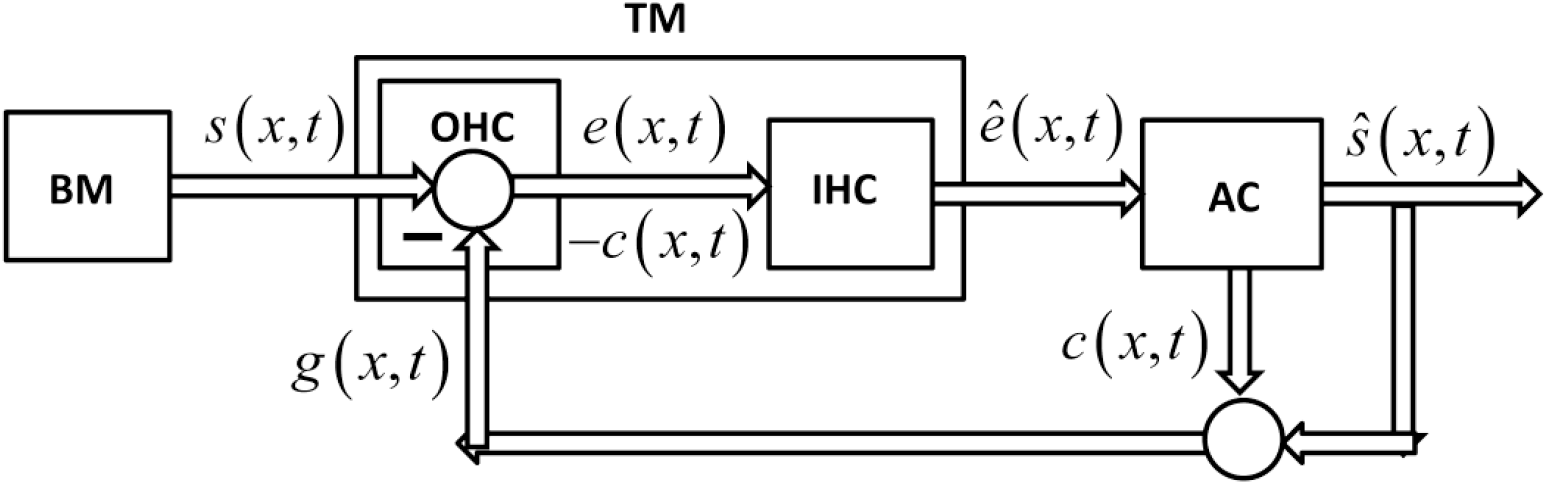
The transduction control system, BM – basilar membrane, TM – temporal membrane, AC –auditory cortex, OHC – outer hair cells, IHC – inner hair cells

The conventional structure of the filter should contain the input comparator of the incoming signal *s*(*x, t*) and the output signal *g*(*x, t*) (the TM vibrations evoked by the activity of efferent fibers of the cochlear nerve). The output signal is supposed to be the sum of two components: 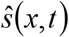 - the best estimate of the incoming signal and *c*(*x, t*) - the low frequency control signal to change the tunnel heights. 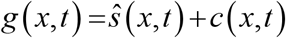. The output of OHC comparator *q*(*x, t*) = *s*(*x, t*)−*g*(*x, t*) contains the sum of the estimation error 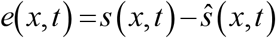 and the control signal *c*(*x, t*), both physically presented as mechanical vibrations and deflections in TM. The estimation error *e*(*x, t*) is amplified to 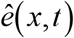 in the molecular IHC filter and forwarded to the auditory cortex. All signals have two dimensional structure (field) being distributed over space and time, so at the level of the auditory cortex the feedback control filter is able to realize all complex processing tasks like filtering in space and time, detection and parameter estimation of any sound wave shape. In scope of this article we will not consider the advanced properties of the filter, concentrating only on its physical implementation in the space of the cochlea. The result of the combined actions is the vibration field propagating on the surface of the TM. The final goal of the space-time filtering is to ensure the minimal amplitude traveling waves along the IHC tunnel. In the ideal case there will be no vibrations observed in the IHC stereocilia tunnel, in spite that in the same time in the BM the amplitudes of traveling waves can be essential. Hence, the information about the sound pressure in the cochlea is taken by the auditory system only from the output of IHC and observed in the output of the feedback loop, i.e. the efferent fibers activity of the cochlea nerve.

## Discussion

The idea to consider TM as the essential part of the cochlea transduction exists in a variety of published articles [23-25]. Some of them also considered the traveling waves propagated in the radial direction in TM [26]. We generalized this idea to the space-time filtering concept, hypothesizing the existence of the molecular filter, which is able to evaluate the traveling wave parameters from the endolymph molecule orientation in the close vicinity of the stereocilia tops.

This allows us to avoid the consideration of mechanical bending of stereocilia of any kind and the mechanical manipulation with the ion channels. The model of the cochlear transduction presented here contains a number of assumptions. The main of them is the weak sound intensity, which leads to the linearization of NS equations and is equivalent to consideration of the tunnel flow akin to the shallow waves [14]. According to the obtained results the IHC are more affected by the back flow in the TM wave propagation, as it is closer to the IHC tops. The longitudinal component of the tunnel flow is declined from the Couette flow shape, but will approach it as far as the amplitude of the TM wave is getting weaker. The new developed concept of the molecular filters built on the tops of IHC let us explain in the more natural way many well known observations, like: high sensitivity, automatic gain control, high precision in TM wave shape evaluation. In spite that the polarization phenomenon of the endolymph molecules was studied early [9,10] in the movement conditions, its implementation for the sensory transduction was discussed only in the vestibular system [20]. There were some attempts to explain the active control properties of OHC by considering them as the elements of the phasing antenna lattice [27]. However, the phasing property of the antenna approach was found to be not efficient, because of small distances between OHC and IHC rows as compared with the relatively long sound wave lengths. Like in the most common control system, the precise estimation of the TM wave shape is supposed to be implemented by the compensation method, where OHC rows are considered to act as the comparator, delivering the error estimation to the IHC molecular filter.

In contrast with the widely discussed active amplifier properties of OHC, the feedback control loop concept presumes that the best estimation is achieved in the most stabilized state of TM, when minimal vibrations are observed and detected by IHC molecular filters. So we don’t need to amplify the BM vibrations, but only to compensate them. The space-time filter is supposed to be self adjusted in a sense to increase its gain for the weak sound signals. For this purpose, the efferent signals from the auditory cortex should contain, in addition to the regular fast component, the slow component intended specifically to slow elongation or shortening of OHC, what in turn, changes the tunnel height. In spite that the exact physical origin of such signal is unclear, there are some evidences that the medial olivocochlear system efferents can modulate the auditory nerve activity on the slow time scale (10-100s) [28]. Recently it was also shown [29] that the changes in the static length of OHC can serve amplifying the magnitude of vibrations of the reticular lamina, what results in increasing of the IHC sensitivity. Increasing or decreasing of the heights results in changes in strength of the endolymph molecules rotation in the IHC molecular filters, and, hence, leads to changes in the probability of IHC excitation. It can be predicted that in the regions of the extremely weak sounds, when forward and backward tunnel flow parts are approximately equal, the control system will try to estimate both parts, what could results in instability (otoacoustic emission), or an appearance of the doubled estimated frequency. The active amplification of BM traveling wave by the vibrations of OHC, which is widely observed and discussed [5,6], is still present, but has another explanation. The elongation and the shortening of the stereocilia bundles in OHC are always reciprocal and perform their action in the opposite direction for TM and BM. Increasing of the BM amplitude by the elongation of the OHC stereocilia provokes the reducing of the TM total amplitude. In reality, the observed increased sensitivity of the cochlea transduction with OHC motility could be explained by better suppression of the BM induced vibrations in the IHC tunnel.

## Conclusion

The new concept of the molecular filters helps explaining the cochlear transduction in the more natural way avoiding the use of the mechanical bending of the inner and outer hair cells and the presence of mechanical tip links for the cell excitation (at least in the regions of the low level sound intensities). The OHC and IHC structures can be considered as a space comparator of two kind of vibrations, directly transferred from BM and evoked from the auditory cortex efferent signals. The molecular filters built on IHC structures, OHC structures and TM can be considered as the parts of the general space-time feedback filter, which performs simultaneously in two roles: evaluation of the BM traveling wave shapes using the compensation method and the automatic gain control of the compensation errors.

